# The depletion mechanism can actuate bacterial aggregation by self-produced exopolysaccharides and determine species distribution and composition in bacterial aggregates

**DOI:** 10.1101/2021.05.11.443568

**Authors:** Patrick R. Secor, Lia A. Michaels, DeAnna C. Bublitz, Laura K. Jennings, Pradeep K. Singh

## Abstract

Bacteria causing chronic infections are often found in cell aggregates suspended in polymer secretions, and aggregation may be a factor in infection persistence. One aggregation mechanism, called depletion aggregation, is driven by physical forces between bacteria and polymers. Here we investigated whether the depletion mechanism can actuate the aggregating effects of *P. aeruginosa* exopolysaccharides for suspended (i.e. not surface attached) bacteria, and how depletion affects bacterial inter-species interactions. We found cells overexpressing the exopolysaccharides Pel and Psl, but not alginate remained aggregated after depletion-mediating conditions were reversed. In co-culture, depletion aggregation had contrasting effects on *P. aeruginosa’s* interactions with coccus- and rod-shaped bacteria. Depletion caused *S. aureus* (cocci) and *P. aeruginosa* (rods) to segregate from each other, *S. aureus* to resist secreted *P. aeruginosa* antimicrobial factors, and the species to co-exist. In contrast, depletion aggregation caused *P. aeruginosa* and *Burkholderia sp*. to intermix, enhancing type VI secretion inhibition of *Burkholderia* by *P. aeruginosa*, leading to *P. aeruginosa* dominance. These results show that in addition to being a primary cause of aggregation in polymer-rich suspensions, physical forces inherent to the depletion mechanism can actuate the aggregating effects of self-produced exopolysaccharides and determine species distribution and composition of bacterial communities.

## Introduction

At sites of chronic infection, bacteria are often found within cell aggregates suspended in polymer-rich host secretions such as mucus, pus, sputum and others (1–3). Aggregated growth is thought important because it can increase the ability of bacteria to survive environmental stresses such as pH and osmotic extremes, as well as host-derived and pharmaceutical antimicrobials (4, 5). Bacterial aggregation also affects disease-relevant phenotypes such as bacterial invasiveness, virulence factor production, and resistance to phagocytic uptake (6–10).

Bacteria can aggregate via bridging aggregation, which occurs when adhesions, polymers, or other molecules bind cells to one another. Another general yet underappreciated mechanism is depletion aggregation (11). Depletion aggregation occurs in environments containing high concentrations of non-adsorbing polymers (12, 13). Such conditions exist in the cytoplasm of eukaryotic cells (11), cystic fibrosis airways (14), wounds (15), biofilm matrices (16), and others settings. Depletion aggregation is initiated when bacteria spontaneously come into close contact with each other (**Fig 1A**), causing the polymers in between cells to become restricted in their configurational freedom, and thus decreasing their entropy. When polymers spontaneously move out from in between bacterial cells (17) a polymer concentration gradient is established across adjacent bacterial cells, producing an osmotic imbalance (i.e., the depletion force) that physically holds the aggregate together (**Fig 1B and C**) (18).

**Figure 1.**
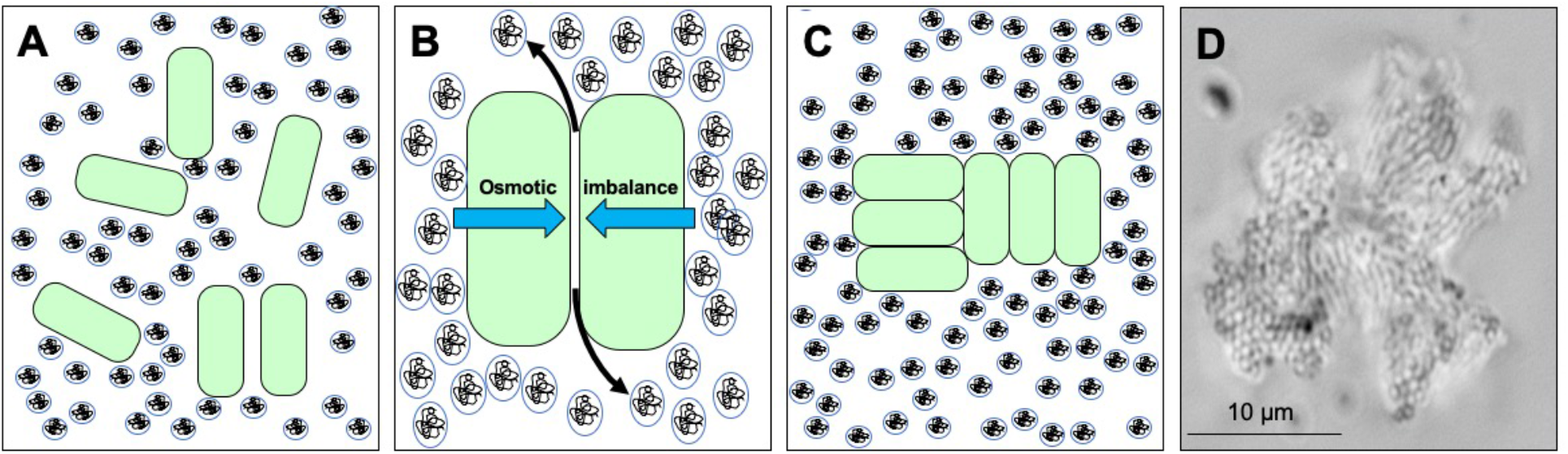
Depletion aggregation is an entropic mechanism that operates to aggregate bacterial cells in environments crowded with non-adsorbing polymers. **(A)** Bacterial cells (green) are suspended in an environment with high concentrations of non-adsorbing polymer (circles). **(B)** Polymers in between cells are restricted in their conformational freedom and spontaneously move out from in between cells (black arrows), increasing their entropy. The polymer concentration gradient across the cells produces an osmotic imbalance (blue arrows). **(C)** The osmotic imbalance (i.e., the depletion force) physically holds the cells together in aggregates. **(D)** Representative image of a *P. aeruginosa* PAO1 depletion aggregate with PEG 35 kDa as the polymer.

While definitions and terminology can vary among investigators, biofilm formation and depletion aggregation can be differentiated by two factors. First, biofilms are generally considered a phenomenon of surface-attached bacteria (19–23), whereas depletion aggregation operates on cells suspended in polymer solutions. Second, biofilm formation is driven by bacterial activity (19, 21, 23) whereas depletion aggregation is a consequence of physical forces generated when high concentrations of polymers are present. If bacteria and polymer concentrations are high enough, aggregation via depletion will occur as default and obligatory outcome unless mechanisms like mechanical disruption or bacterial motility produce stronger counteracting forces. The dependency on environmental conditions also means that a reduction in polymer concentration will cause aggregate to disperse, unless other mechanism of bacterial adhesion supervene.

Previous work has shown that the concentrations host-derived polymers like mucin, DNA, and F-actin found at infection sites cause bacterial depletion aggregation (as do model polymers like PEG), that and that depletion aggregation causes bacteria an antibiotic-tolerant phenotype (14). Here we investigated how the depletion mechanism affects aggregation mediated by *P. aeruginosa*’s biofilm exopolysaccharides and recently identified in cystic fibrosis sputum. We also investigated how depletion aggregation affects the interactions between bacterial species that may co-exist *in vivo*.

## Results

### Depletion aggregation can actuate bridging interactions by exopolysaccharides

Biofilm formation is generally thought to occur when surface-attached cells accumulate by growth, moving towards each other, or recruitment from overlying media; and produce exopolysaccharides and other matrix components that enable them to stick together via bridging interactions (24, 25). Unlike surface-attached cells, bacteria suspended in solutions are subject to random (i.e. Brownian) movement or fluid flows that can disperse them, reducing cell to cell contact and the potential for bridging interactions. These points led us to hypothesize that in addition to being a primary aggregation mechanism, the depletion mechanism could facilitate bridging interactions mediated by biofilm matrix components.

*P. aeruginosa* encodes three exopolysaccharides. Pel is a cationic polymer composed of partially acetylated N-acetylgalactosamine and N-acetylglucosamine (26), Psl is a neutral polymer containing glucose, mannose, and rhamnose (27), and alginate is a negatively-charged polymer composed of mannuronic and guluronic acid (28, 29). We first tested wild-type *P. aeruginosa* that are capable of producing all of these three exopolysaccharides (30, 31). As seen previously, wild type *P. aeruginosa* exposed to the model polymer PEG 35 kDa rapidly aggregated via the depletion mechanism, and immediately diluting the polymer by adding PBS caused the aggregates to disperse whereas adding additional PEG did not. As noted above, reversibility with dilution is a hallmark of depletion aggregation, as it is driven by crowding effects of environmental polymers.

We reasoned that longer aggregation time periods could enable wild type *P. aeruginosa* to produce biofilm exopolysaccharides and adhesive bridging interactions. However, disaggregation of wild-type *P. aeruginosa* was noted even in aggregates that were held together by polymer exposure for 18 hrs (**Fig 2A; Movie 1**). To determine if high level expression of exopolysaccharides could cause depletion-induced aggregates to persist after polymer dilution we repeated these experiments using *P. aeruginosa* overproducing alginate (due to a mutation in alginate regulator, *mucA*) and Pel and Psl (due to induced expression from a P_BAD_ promoter). *P. aeruginosa* over-expressing Pel and Psl remained aggregated after PBS (or PEG) dilution (**Fig 2B and C**), whereas the strain over producing alginate did not (**Fig 2D**).

**Figure 2.**
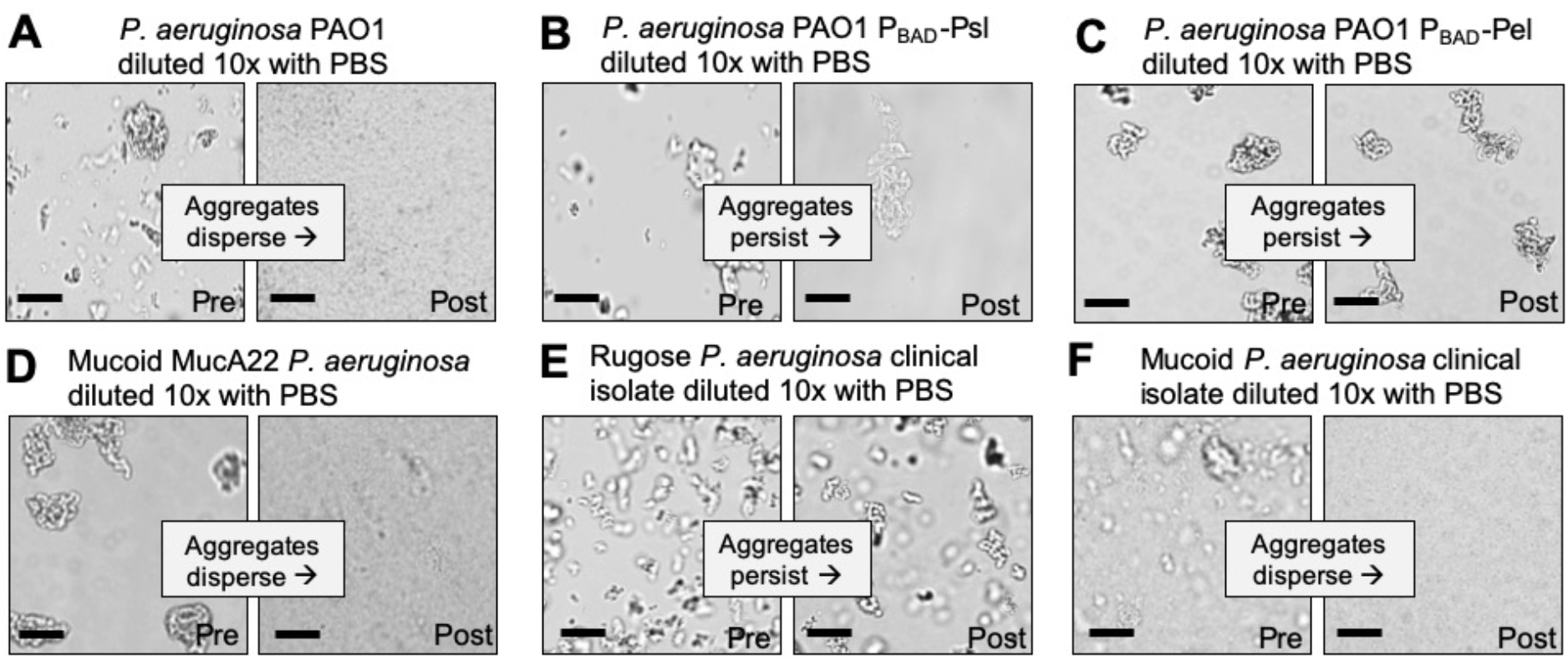
Depletion aggregate dispersal phenotypes of *P. aeruginosa* laboratory strains and CF clinical isolates. **(A-F)** Aggregate dispersal of the indicated strain and isolates was measured. Depletion aggregation was induced with 30% w/vol PEG 35 kDa for 18 hours. Depletion aggregates were then diluted 10X with PBS and representative images were acquired immediately pre- and immediately post-dilution. See also Figure S1 and Movie S1. Scale bar 40 μm.

We also studied clinical isolates taken from cystic fibrosis patients (32) that are known to overexpress Pel, Psl, or alginate. All 10 *P. aeruginosa* CF clinical isolates tested that over-produced the exopolysaccharides Psl or Pel (6, 33) formed dilution-resistant depletion aggregates (**Fig 2E, Table 1**), consistent with observations in corresponding engineered lab strains. In contrast, all (9/9) alginate-overproducing clinical isolates (i.e. mucoid strains) had a reversible aggregation phenotype (**Fig 2F, Table 1**), consistent with observations with the mucoid PAO1 *mucA22*. Collectively, these results indicate that under the conditions tested, Pel and Psl can stabilize aggregates formed by the depletion mechanism if they are highly expressed while alginate does not. The different chemical compositions or other physical properties such as charge may explain differences in aggregate reversibility.

**Table 1.** *P. aeruginosa* morphology and aggregate reversibility phenotypes.

| Strain | Morphology | Reversible aggregation? |
| --- | --- | --- |
| PAO1 | Non-mucoid | Yes |
| PAO1 $\Delta wspF/pslD$;<br>pBAD::Pel | Non-mucoid | No |
| PAO1 $\Delta wspF/pelF$;<br>pBAD::Psl | Non-mucoid | No |
| PDO300 mucA22 | Mucoid | Yes |
| PAO1 $\Delta mucA$ | Mucoid | Yes |
| Clinical Isolate 2-6.3 | Mucoid | Yes |
| Clinical Isolate 29-14 | Mucoid | Yes |
| Clinical Isolate 7-15.4 | Mucoid | Yes |
| Clinical Isolate 9-19.6A | Mucoid | Yes |
| Clinical Isolate W1 | Mucoid | Yes |
| Clinical Isolate W2 | Mucoid | Yes |
| Clinical Isolate W3 | Mucoid | Yes |
| Clinical Isolate W4 | Mucoid | Yes |
| Clinical Isolate W5 | Mucoid | Yes |
| Clinical Isolate 27-6.4 | Rugose | No |
| Clinical Isolate 28-17.9 | Rugose | No |
| Clinical Isolates 29-5.6 | Rugose | No |
| Clinical Isolate 14-4.2 | Rugose | No |
| Clinical Isolate 17-6.6 | Rugose | No |
| Clinical Isolate S1 | Rugose | No |
| Clinical Isolate S2 | Rugose | No |
| Clinical Isolate S3 | Rugose | No |
| Clinical Isolate S4 | Rugose | No |
| Clinical Isolate S5 | Rugose | No |

### Cell shape determines species distribution in depletion aggregates

Theory predicts that bacteria aggregated by the depletion mechanism will be arranged to minimize the amount of volume occupied, as efficient packing will increase the space available for polymers and concomitant entropy gains. This effect should cause bacteria with similar shapes to be arranged together, and bacterial with different shapes to separated, unless bacterial activity intervenes. To test this hypothesis, we mixed *P. aeruginosa, Burkholderia cenocepacia, Escherichia coli* (rods) and *Staphylococcus aureus* (a coccus) bearing different florescent labels in various combinations in PEG 35 kDa, and examined species distribution by microscopy.

Polymer-mediated depletion aggregation caused cocci shaped species (*S. aureus*) to segregate from rods (*P. aeruginosa* and *B. cenocepacia*). In some cases, entire aggregates appeared composed of single species. In other cases, sections of mixed-species aggregates were composed primarily of either the rod or cocci-shaped species (**Fig 3A and B**). In contrast, depletion aggregation caused bacteria with similar cell shapes (i.e. differentially labeled *P. aeruginosa* with *P. aeruginosa, or P. aeruginosa* with *E. coli*) to intermix (**Fig 3C and D**). Similar results were seen using mixtures of formalin-killed *P. aeruginosa* and *S. aureus*, and formalin-killed *P. aeruginosa* and 2 μm diameter spherical beads similarly sized as *S. aureus* **(Fig S2A and B)**. These experiments, along with previous work using inert particles (34), show that physical forces mediating depletion aggregation cause like-shaped bacteria to intermix, and differently shaped bacteria to separate. The physical arrangement of bacterial species in aggregates can affect competitive and cooperative interactions (see below).

**Figure 3.**
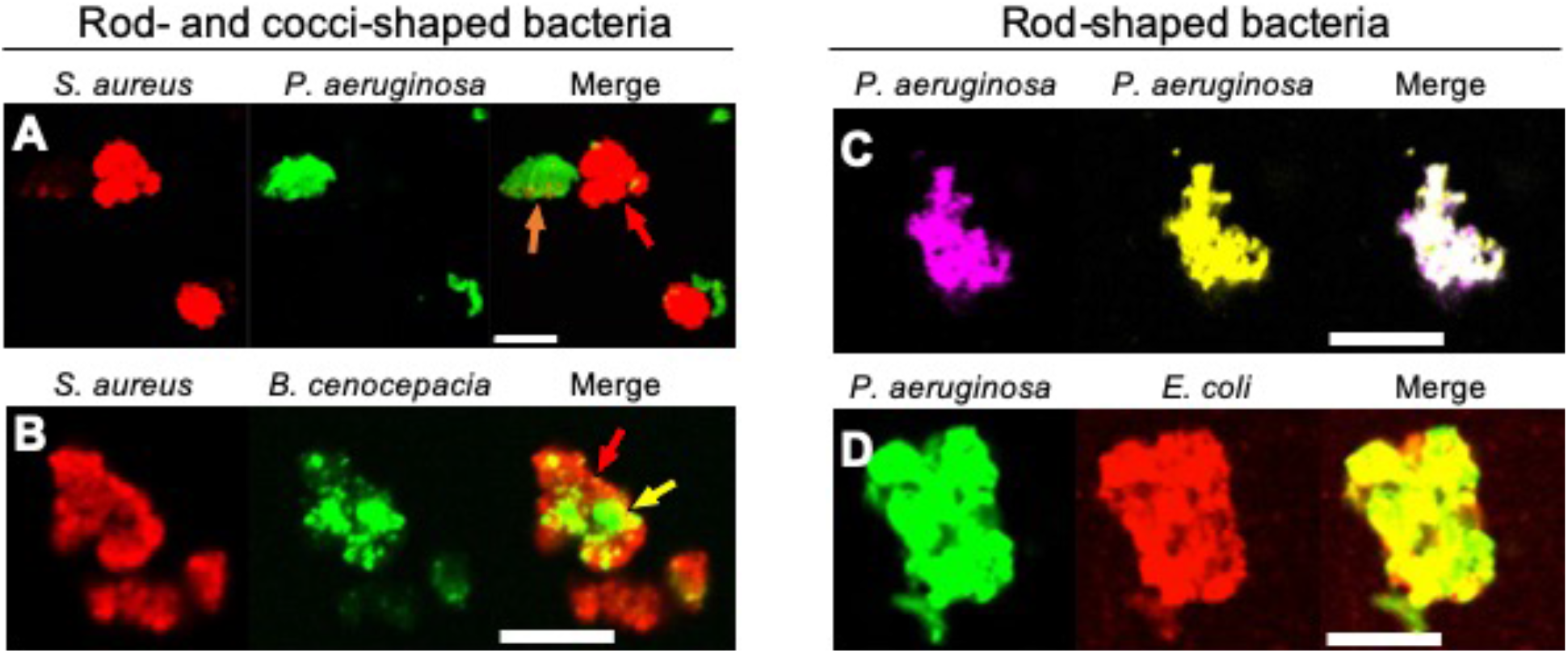
Depletion aggregation spontaneously segregates bacteria with different cell shapes. Equal numbers of the indicated species were mixed prior to the addition of PEG 35 kDa to induce depletion aggregation. Aggregates were imaged 18-h later. Combinations of rod- and cocci-shaped bacteria are shown in **(A and B)** and combinations of rod-shaped bacteria are shown in **(C and D)**. Scale bar 30 μm.

### Depletion aggregation promotes antimicrobial tolerance in *S. aureus*

Our finding that depletion aggregations can determine the physical arrangement of species led us to investigate its effects on interspecies interactions. *P. aeruginosa* and *S. aureus* are often co-isolated from CF airways (35, 36) and wounds (37, 38) for long durations. However, in laboratory co-cultures, *P. aeruginosa* rapidly inhibits *S. aureus* by quorum-regulated antimicrobials such as rhamnolipids, hydrogen cyanide, phenazines, quinolones, and others (39–43). Because aggregation can increase antimicrobial tolerance (44, 45), we hypothesized that depletion aggregation could enhance the ability of *S. aureus* to co-exist with *P. aeruginosa*.

Similar to previous studies, (39–43) we found that wild-type *P. aeruginosa* severely inhibited *S. aureus* in non-aggregated broth co-cultures (**Fig 4A**), and inhibition was diminished if *P. aeruginosa’s* main quorum sensing systems were genetically inactivated (i.e. Δ*lasR/rhlR* PAO1; p<0.01) (**Fig 4A**, compare white bars). However, in co-cultures exposed to PEG 35 kDa to induce depletion aggregation, wild-type *P. aeruginosa* killing of *S. aureus* was reduced by over 10-fold (**Fig 4A**, black bars).

**Figure 4.**
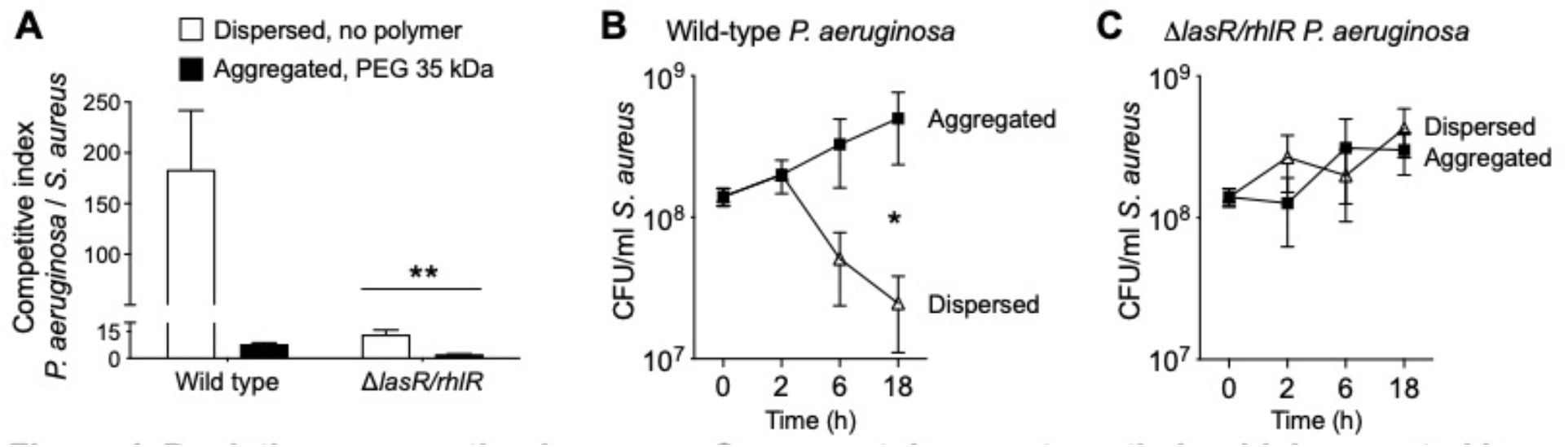
Depletion aggregation increases *S. aureus* tolerance to antimicrobials secreted by *P. aeruginosa*. **(A)** Equal numbers (10^7^ CFUs) of *S. aureus* and *P. aeruginosa* (wild-type PAO1 or Δ*lasR/rhlR*) were cocultured in LB supplemented with 30% w/vol PEG 35 kDa where indicated. After 18-h, viable bacteria were enumerated by serial dilution and plating and plotting the competitive index (change [final/initial] in *P. aeruginosa* vs. *S. aureus* CFUs). Results are the mean ± SD, N=3 for each condition; **p<0.01 relative to wild type. **(B and C)** *S. aureus* (10^8^ CFU/ml) was added to filter sterilized supernatants collected from wild-type or Δ*lasR/rhlR P. aeruginosa* overnight cultures supplemented with 30% w/vol PEG 35 kDa where indicated. Viable *S. aureus* was enumerated by serial dilution and plating at the indicated times. Results are the mean ± SD, N=3 for each condition and timepoint; *p<0.02.

Our previous finding that depletion aggregation caused marked antibiotic tolerance in *P. aeruginosa* (14) led us to hypothesize that depletion-mediated tolerance explained *P. aeruginosa-S. aureus* co-existence in aggregates. We tested this by exposing dispersed and depletion-aggregated *S. aureus* to filter-sterilized culture *P. aeruginosa* supernatant found that dispersed *S. aureus* were ~10-fold more sensitive to killing after 18 hrs (**Fig 4B**). Control experiments indicate that PEG did not diminish the antimicrobial activity of *P. aeruginosa* supernatants (see **Fig S3**), and that the inhibitory effects were mediated by quorum-controlled factors **(Fig 4C)**. These results indicate that depletion aggregation can promote co-existence of *P. aeruginosa* and *S. aureus* by enhancing *S. aureus* tolerance to quorum-controlled antimicrobials secreted by *P. aeruginosa*.

### Depletion aggregation promotes contact-dependent bacterial antagonism

In addition to secreted factors, *P. aeruginosa* and other bacteria also possess competitive mechanisms that depend upon cell-to-cell contact. One mechanism is type VI secretion (TSS) in which a needle-like apparatus delivers toxins into neighboring cells (46). Our finding that depletion aggregation causes like-shaped bacterial cells intermix in aggregates led us to hypothesize that it would promote TSS-mediated bacterial antagonism.

To test this, we mixed *P. aeruginosa* which capable of T6SS with *Burkholderia thailandensis*, a TSS-susceptible rod-shaped Gram-negative bacterium (47). In dispersed conditions, no *P. aeruginosa*-*B. thailandensis* antagonism was apparent over 24 hours, as the ratio *P. aeruginosa* to *B. thailandensis* remained unchanged (**Fig 5A**). In contrast, *P. aeruginosa* outcompeted *B. thailandensis* in depletion aggregates as measured by viable counts (**Fig 5A**) and visually assessing differentially-labeled species (**Fig 5B)**. Notably the competitive advantage of *P. aeruginosa* was eliminated by genetically inactivating TSS (i.e. PAO1 *ΔclpV1* (**Fig 5C**)). Taken together, these results demonstrate that depletion aggregation can facilitate contact-dependent mechanisms of bacterial antagonism.

**Figure 5.**
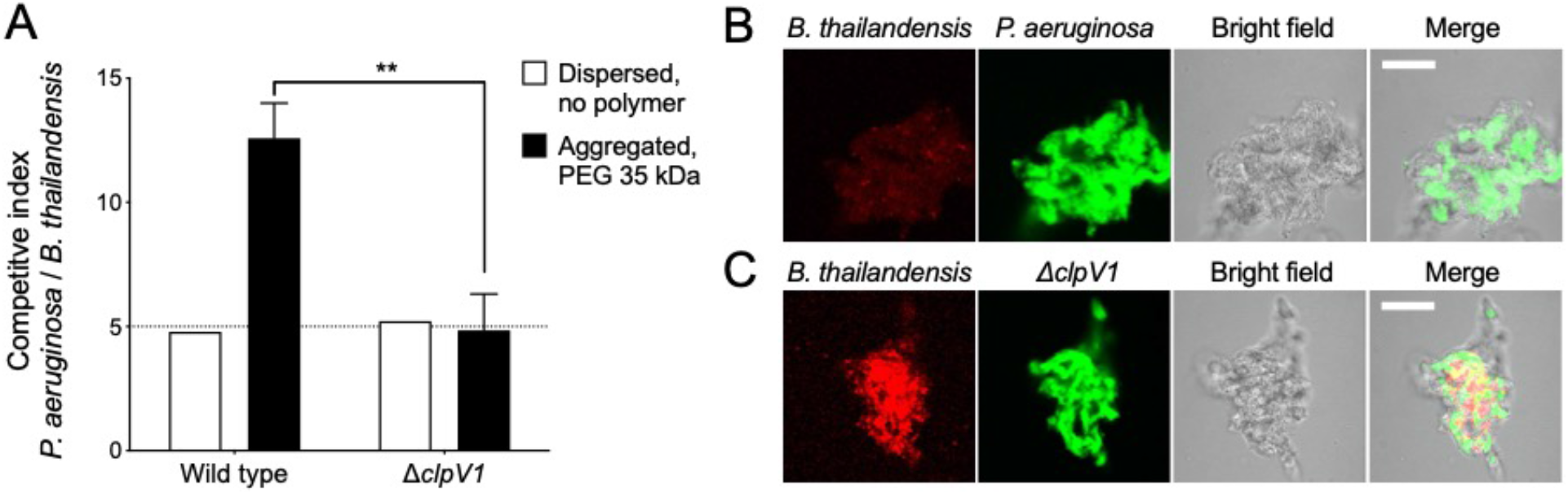
Depletion aggregation promotes contact-dependent bacterial competition. **(A)** The outcome of competitions between *B. thailandensis* and either wild-type or Δ*clpV1 P. aeruginosa* are shown. Initial cultures contained 1×10^8^ CFU/ml *P. aeruginosa* and 2×10^7^ CFU/ml *B. thailandensis*. Results are after 24-h of co-culture in the indicated conditions and are the mean ± SD, N=3 for each condition; **p<0.01. **(B and C)** Fluorescent microscopy was used to visualize depletion aggregates after 24-h of co-culture with 30% PEG 35 kDa. Representative images are shown with *B. thailandensis strains* in red and *P. aeruginosa* in green. Scale bar, 30 μm.

## Discussion

Previous observations by a number of groups has shown that pathogens causing chronic infection like those in cystic fibrosis and wounds are generally found to be living in aggregates suspended in polymer-rich secretions, rather than as surface attached biofilms (1, 2, 10, 38, 48–56). Our previous work shows that physical forces produced by polymers found at infection sites can cause bacteria to form suspended aggregates by the depletion mechanism, and depletion aggregation produces disease-relevant phenotypes (14). In this study we found that depletion aggregation can actuate bridging interactions mediated by two of *P. aeruginosa*’s self-produced biofilm polysaccharides; cause bacteria with like shapes to arrange together and bacteria with different shapes to segregate, and has different effects on bacterial competition mechanisms mediated by secreted factors and cell-to-cell contact.

Surface attachment is thought to be fundamental to biofilm formation; sensing and adhering to surfaces induces physiological responses important in biofilm growth, and attachment keeps nascent biofilm-forming cells from dispersing (from random movement or fluid flows) before the matrix binds them together (23). Our previous work and current experiments raise the possibility that the depletion mechanism might serve somewhat similar functions as attachment surfaces. Previously we found that like surface attachment (57), depletion aggregation can induce stress responses in *P. aeruginosa* that mediate antibiotic tolerance (14). Our current experiments show that depletion aggregation also brings suspended cells together and can promote adhesion by self-produced polymers. One important caveat is that in the conditions used here, exopolysaccharide overexpression was required as *P. aeruginosa* PAO1 capable of producing “wild-type” levels of polysaccharides did not exhibit matrix-mediated adhesion even after long periods of depletion aggregation. Notably, mutant strains constitutively expressing EPS can be isolated from infected CF subjects (58), and *in vivo* conditions could induce expression of matrix polysaccharides to levels needed to cause bridging aggregation.

Our findings also have implications for interspecies interactions that may occur in infections. The experiments showing that depletion aggregation increases tolerance of *S. aureus* to antimicrobials produced by *P. aeruginosa* (**Fig 6A**) could help explain how *P. aeruginosa* and *S. aureus* can co-exist in chronic infections like wounds and CF lungs, but are difficult to maintain in liquid co-cultures. While the underlying mechanism remains to be characterized, our previous work showing that that depletion aggregation induces the SOS stress response (14) raises the possibility that a similar phenomenon operates in *S. aureus* (59, 60). If general stresses were induced, aggregated *S. aureus* may exhibit tolerance to other disease-important stresses including antibiotics.

**Figure 6.**
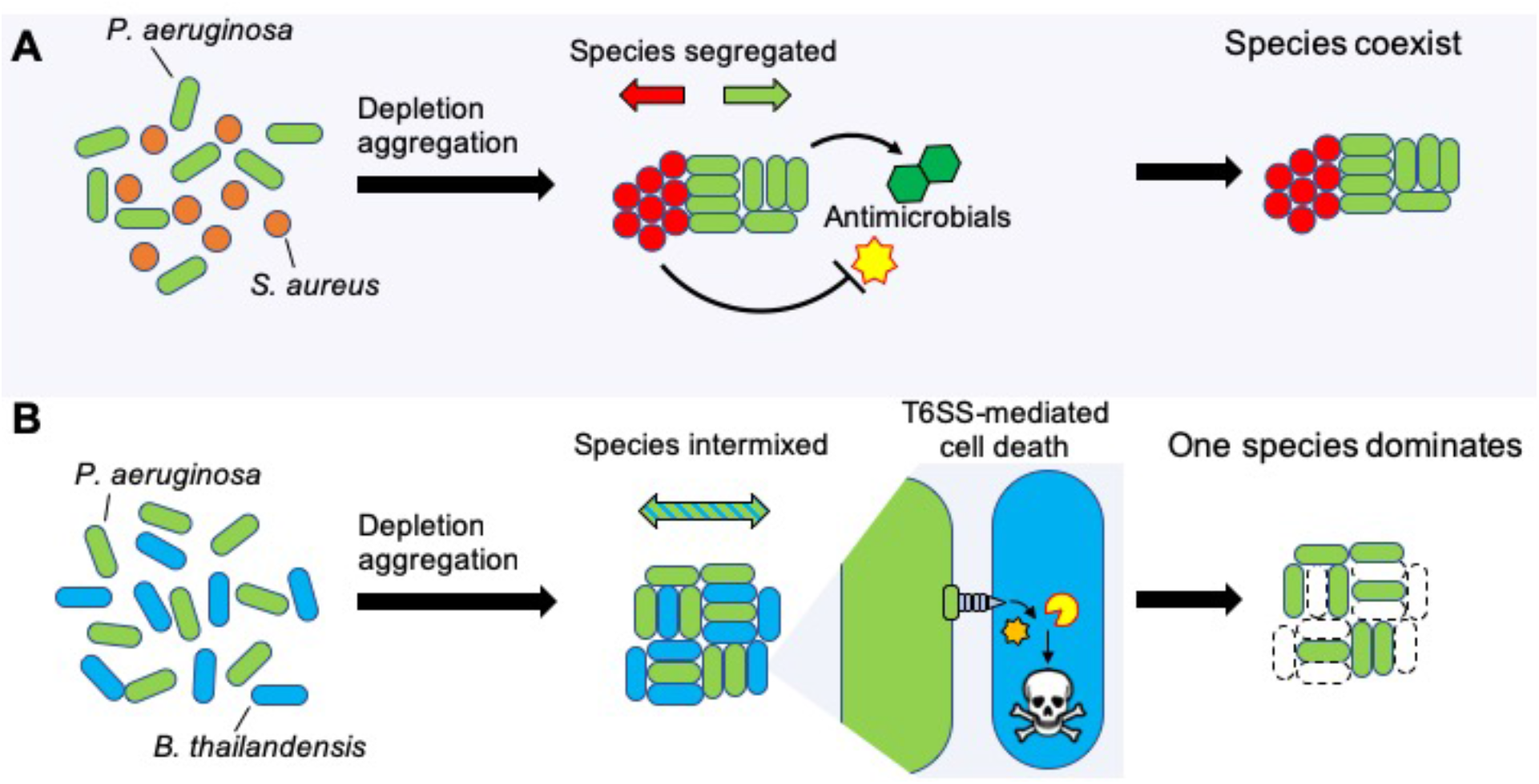
Model depicting how depletion aggregation affects bacterial competition and species distribution in aggregates. **(A)** Depletion aggregation causes bacteria with different cell shapes to spontaneously segregate. When *P. aeruginosa* and *S. aureus* were co-cultured under conditions promoting depletion aggregation, *S. aureus* tolerated antimicrobials secreted by *P. aeruginosa*, promoting species coexistence. **(B)** When two rod-shaped species such as *P. aeruginosa* and *B. thailandensis* are aggregated by the depletion mechanism, species segregation is not observed and contact-dependent T6SS-mediated killing is promoted, allowing *P. aeruginosa* to dominate.

The effect of depletion aggregation to intermix species with similar shapes, and segregate species dissimilar shapes could have wide ranging effects. One consequence we demonstrated is enhanced efficacy of TSS-mediated inhibition of rod shaped *Burkholderia sp*. by rod-shaped *P. aeruginosa*, as TSS is dependent upon species intermixing and cell-to-cell contact (**Fig 6B**). Such interactions could contribute to the ability of *P. aeruginosa* to dominate other rod-shaped pathogens such as *Haemophilus influenzae* and *Stenotrophomonas maltophilia* (35, 36, 61–63) in CF airways. Depletion aggregation could likewise enhance or inhibit other close-range mechanisms that depend on contact or have short diffusion distances (like oxidants) depending on whether species are of similar or dissimilar shapes. In addition, in settings where depletion aggregation is maintained for long durations (i.e. polymers are continuously present), its effects on species arrangement could shape co-evolutionary trajectories of species, as the within-aggerate arrangement of cells likely affects selection, competition, and cell migration.

Our study had several limitations. For example, we used a non-biological polymer (PEG) at a specific concentration (30% w/vol) with a defined molecular weight (PEG 35 kDa) to induce depletion aggregation as use of a defined polymer limited variability and the transparency of PEG enhanced microscopy. While it is possible that biological polymers could produce different results, our previous work shows that depletion aggregation by DNA and mucin at concentrations found at infection sites cause similar aggregate morphology and antibiotic tolerance phenotypes as PEG (14). We also recognize that varying polymer size and molecular weight will affect the strength of the aggregating force, and these variables were not examined here. An additional limitation was that our experiments used laboratory strains and a handful of *P. aeruginosa* clinical isolates. Clinical isolates with different biological characteristics could affect depletion-mediated bacteria-bacteria interactions. For example, LPS or other cell envelope modifications that arise *in vivo* (62) could change cell surface charge or hydrophobicity, which could affect depletion-mediated bacteria-bacteria or bacteria-polymer interactions.

Much research in model systems has been devoted to understanding bacterial sensing and signaling pathways, purpose-evolved genetic programs, and quasi-social cooperation that shape bacterial phenotypes important in chronic infections. The data presented here show that basic thermodynamic forces inherent to polymer-rich environments can have marked effects on complex bacterial behaviors including aggregation, stress survival, and interspecies competition. New strategies to manipulate pathogenesis phenotypes will require understanding the relative contributions of bacterially-driven processes and mechanisms caused by physical forces in the environment. Generating such knowledge is challenging as cause-and-effect relationships are difficult to discern thorough the observational studies possible with human samples, and because animal models representing chronic infection have been difficult to develop. Ultimately, understanding may come from studying the effects of interventions that manipulate bacterially-driven processes or the physical environment present at infection sites.

## Acknowledgments

We are grateful to Joseph Mougous for sharing the *clpV1* mutant and *Burkholderia* strains.

## Methods

### Chemicals/growth media/strains

Growth media (Lysogeny broth, LB), polyethylene glycol MW 2,000 and 35,000 Da, and antibiotics were purchased from Sigma. Strains and their sources are listed in **Table 2**.

**Table 2.** Strains used in this study.

| Strain | Description | Source |
| --- | --- | --- |
| PAO1 | Wild type | (64) |
| PAO1 $\Delta wspF/pslD$;<br>pBAD::Pel | Deletion of *wspF* and *pslD*; arabinose-inducible Pel operon | (65) |
| PAO1 $\Delta wspF/pelF$;<br>pBAD::Psl | Deletion of *wspF* and *pelF*; arabinose-inducible Psl operon | (26) |
| MucA22 (PDO300) | A *mucA22* allele derivative of PAO1 constructed by allelic exchange | (66) |
| PAO1 $\Delta mucA$ | Contains a truncated *mucA* allele | (67) |
| Clinical Isolates | *P. aeruginosa* clinical isolates from various patients | (32) |
| PAO1 $\Delta lasR/rhlR$ | Deletion of *lasR* and *rhlR* | (68) |
| PAO1 $\Delta clpV1$ | Deletion of *clpV1* | (46) |
| PAO1 attTn7::GFP | Constitutive expression of GFP | (69) |
| PAO1 $\Delta clpV1$; attTn7::GFP | Deletion of *clpV1*; constitutively expressing GFP | (47) |
| PAO1 attTn7::TFP | Constitutive expression of TFP | (20) |
| PAO1 attTn7::YFP | Constitutive expression of YFP | (20) |
| *E. coli* pUCP18-*mCherry* | Carries plasmid expressing IPTG-inducible mCherry | (70) |
| *B. thailandensis* E264 | Wild type | (71) |
| *B. thailandensis* E264<br>attTn7::mCherry | Constitutive expression of mCherry | (47) |
| *B. cenocepacia* K56-2<br>attTn7::GFP | Constitutive expression of GFP | (72) |
| *S. aureus* SH1000 | Wild type | (73) |
| *S. aureus* pCE-SarA-*mCherry* | Constitutive expression of mCherry | (74) |

### PEG-induced depletion aggregation of bacteria

For PEG-induced depletion aggregation, bacteria were added at the indicated densities to either LB diluted 4:6 with distilled water or LB diluted with 50% PEG 35 kDa (w/vol) prepared in distilled water to ensure that nutrient concentrations were the same in dispersed and aggregated conditions. LB was diluted with water or 50% w/vol PEG 35 kDa for all experiments described unless noted otherwise. Cultures were then incubated on a roller (60 rpm) at 37°C unless indicated otherwise.

### Aggregate reversibility assays

The indicated bacterial strains were grown overnight in full-strength LB. One hundred μl of overnight cultures were used to inoculate 3 ml of LB+PEG 35 kDa. After 18-h of growth, 100 μl of the indicated cultures were removed to a 1.5 ml tube containing 900 μl of either 1x PBS or PBS supplemented with 30% w/vol PEG 35 kDa and vortexed. Imaging was performed on 50 μl culture aliquots pre- and post-dilution using a Leica DM1000 LED microscope by spotting onto a glass slide. Aggregate dispersal was scored by eye by comparing to undiluted control cultures.

### Bacterial competition assays

*S. aureus* SH1000 (73) and *P. aeruginosa* PAO1 (64) were grown overnight at 37°C with shaking in LB broth. *S. aureus* and *P. aeruginosa* were pelleted and resuspended at 10^8^ CFU/ml in fresh LB broth. One hundred μl of each culture was added to 2 ml LB supplemented with either 30% w/vol PEG (35 kDa or 2 kDa) where indicated. Bacteria were grown in co-culture for 18 h and viable bacteria were enumerated by serial dilution and plating on LB plates. Colony morphology was used to differentiate *P. aeruginosa* from *S. aureus*.

For experiments investigating the effects of quorum-regulated antimicrobials on *S. aureus* killing, *P. aeruginosa* PAO1 or *ΔlasR/rhlR* (68) were grown overnight at 37°C with shaking in 50 ml LB broth in a 250 ml flask. Bacteria were removed by centrifugation (10 minutes, 9,000 x g) and supernatants were filter sterilized using bottle top vacuum filters with 0.2 μm pore size (Millipore). PEG 2 kDa or 35 kDa was added to these supernatants to a final concentration of 30% w/vol where indicated. *S. aureus* was inoculated into *P. aeruginosa* supernatants at 10^8^ CFU/ml and cultured for 6 h at 37°C on a roller at 60 rpm. Viable *S. aureus* were enumerated by serial dilution and plating onto LB agar plates.

To investigate TSS mediated killing, *P. aeruginosa* PAO1, *ΔclpV1* (46), and *B. thailandensis* E264 (71) were grown overnight at 37°C with shaking in LB broth. Bacteria were resuspended in fresh LB at 10^9^ CFU/ml. One hundred μl containing 1×10^8^ CFU *P. aeruginosa* PAO1 or Δ*clpV1* and 100 μl containing 2.0×10^7^ CFU *B. thailandensis* were added to 800 μl LB or the indicated polymer solutions and incubated in co-culture for 24 h at 37°C on a roller at 60 rpm. Viable bacteria were enumerated by serial dilution and plating on LB plates. Colony morphology was used to differentiate *P. aeruginosa* from *B. thailandensis*. For fluorescent imaging of aggregates, strains PAO1 or *ΔclpV1* constitutively expressing GFP (PAO1 attTn7::*GFP*, (69)) were co-cultured with *B. thailandensis* E264 attTn7::*mCherry* for 24 hours (47). Image analysis is described below.

### Fluorescent microscopy

*S. aureus* SH1000 carrying the fluorescent reporter pCE-SarA-mCherry (74), *P. aeruginosa* PAO1 attTn7::*GFP*, PAO1 attTn7::*TFP* (20), PAO1 attTn7::*YFP* (20), *Escherichia coli* carrying pUCP18-mCherry (70), *B. cenocepacia* K56-2 attTn7::*GFP* (72) and *B. thailandensis* E264 attTn7*::mCherry* were co-cultured as indicated. Depletion aggregates assembled from dead bacteria were prepared by washing and resuspending overnight cultures of PAO1 YFP or PAO1 TFP in PBS at a concentration of 10^9^ CFU/ml. Formaldehyde (16%, Thermo) was added slowly to bacteria while vortexing to a final concentration of 4% vol/vol. Bacteria were allowed to fix for 30 minutes with constant mixing to prevent bacteria from clumping. Cells were then centrifuged for 10 minutes at 9,000 x g, washed twice with PBS, and resuspended in 1 ml PBS. Complete bacterial killing was confirmed by plating fixed bacteria on LB agar. One hundred μl of the indicated fixed strains were added to 2 ml PBS or PBS+30% PEG 35 kDa. Bacteria were incubated in a 37°C in a roller at 60 rpm and visualized at the indicated times using a Zeiss LSM 510 confocal laser-scanning microscope. Image series were processed using Volocity (Improvision).

---

**Figure S1.**
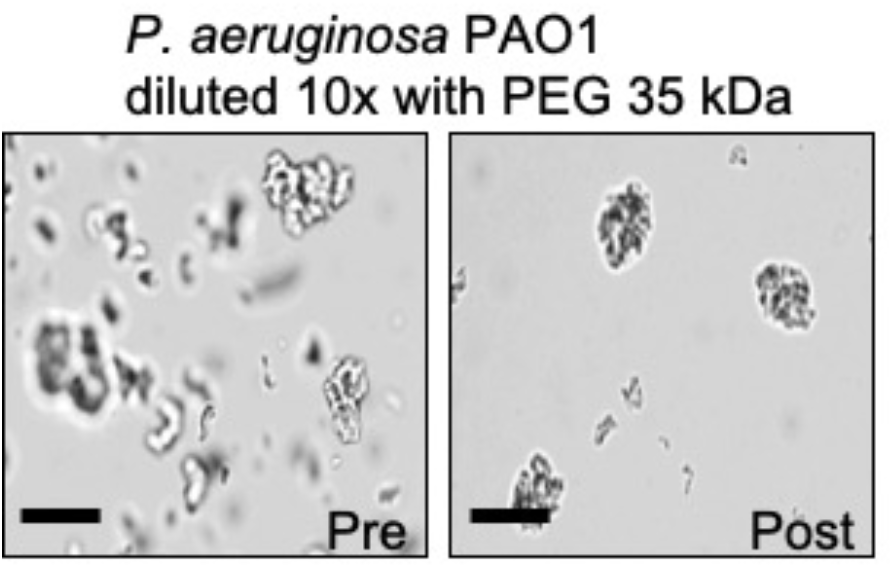
Depletion aggregation was induced with 30% w/vol PEG 35 kDa for 18 hours. *P. aeruginosa* PAO1 depletion aggregates were then diluted 10X with additional PEG 35 kDa. Scale bar 40 μm.

**Figure S2.**
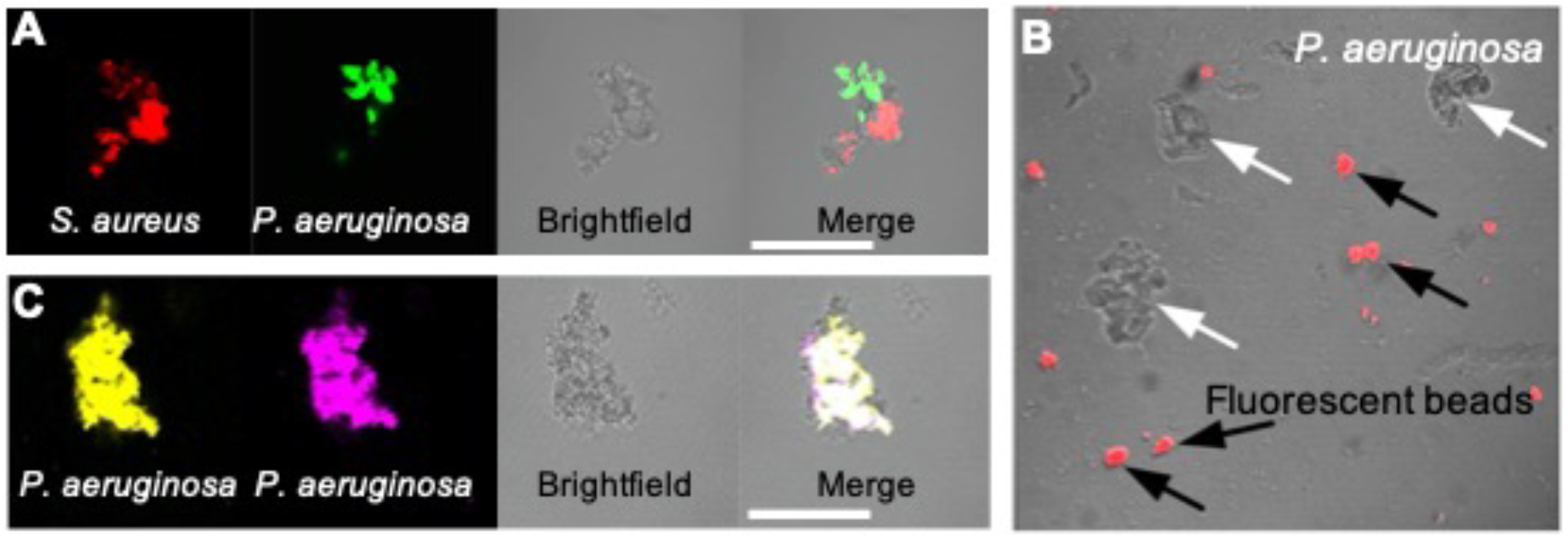
Depletion aggregation operates on dead cells and inert latex beads. **(A and C)** Depletion aggregation was induced with PEG 35 kDa using combinations of dead formalin-fixed cocci and rods. Fluorescent microscopy was used to image aggregates after 18-h of growth. Note species segregation in B (rod+cocci) but not in C (rod+rod). Bar, 30 μm. **(B)** *P. aeruginosa* (rod, white arrows) and fluorescent spherical latex beads (2 μm diameter, black arrows) were aggregated using 30% w/vol PEG 35 kDa for 18-h and imaged using fluorescent and brightfield microscopy. Bar, 30 μm.

**Figure S3.**
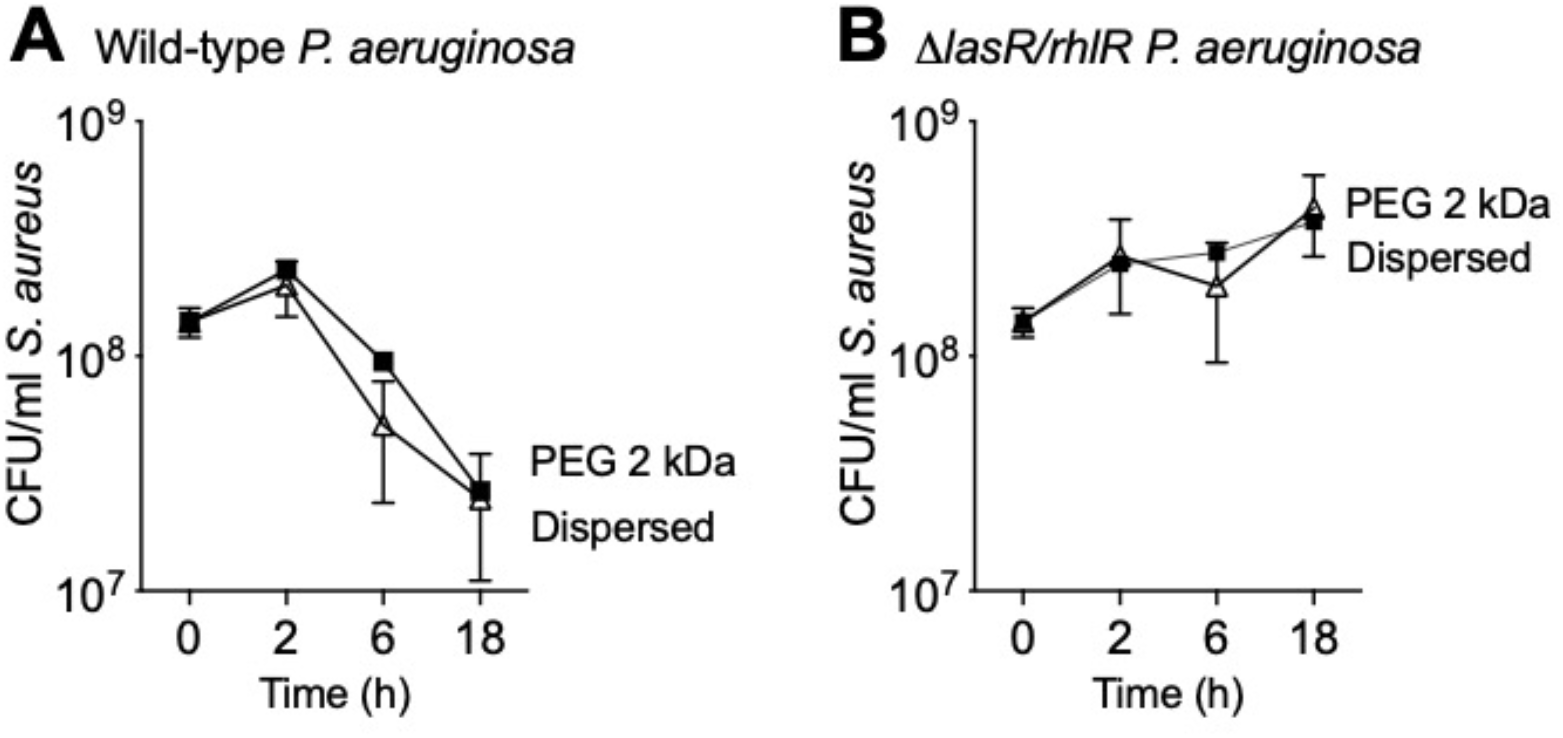
PEG does not inactivate antimicrobials present in *P. aeruginosa* supernatants. One possible explanation for the reduced killing of aggregated *S. aureus* (see Fig 4) was that PEG somehow inactivated antimicrobials present in wild-type *P. aeruginosa* supernatants. To address this possibility, we used a lower molecular weight PEG (PEG 2 kDa). As polymer molecular weight decreases, the polymer concentration required to induce depletion aggregation of a given number of cells increases (12, 14). Thus, PEG 2 kDa does not promote depletion aggregation at 30% w/vol (14). Dissolving PEG 2 kDa into wild-type *P. aeruginosa* supernatants **(A)** did not reduce *S. aureus* inhibition compared to polymer-free controls **(B)**, indicating that PEG did not inactivate antimicrobials present in *P. aeruginosa* supernatants.

